# PARTITIONING AMONG NICHE AND NEUTRAL EXPLANATIONS FOR METACOMMUNITY PATTERNS IN CERRADO STREAM FISH COMMUNITIES

**DOI:** 10.1101/2020.04.06.027417

**Authors:** Thiago Bernardi Vieira, Liriann Chrisley Nascimento da Silva, Lilian Casatti, Renato Romero, Francisco Leonardo Tejerina Garro, Pedro De Podestà Uchôa de Aquino, Paulo dos Santos Pompeu, Paulo De Marco

## Abstract

The Species Sorting concept, one of the models developed to explain patterns in metacommunity structure, suggests that relationships between biological communities and environmental conditions is the basic means of the species selection processes. A second concept is neutral theory, and the idea of neutral dynamics underpinning metacommunity structure, cannot be overlooked. The third mechanism is the Mass Effect concept, that focuses on the interaction between environmental condition and neutral effects. In the present study, we partitioned fish communities in streams between niche and neutral theory concepts, identifying the best representation of metacommunity structure, and assessed if linear and hydrographic distance were equivalent in the representation of neutral processes. The result points to the importance of species sorting mechanisms in structuring fish communities with neutral processes best represented by the linear distances. On the other hand, the best representation of species’ niches was achieved with average values and variance of the local conditions.

## Introduction

Fish stream communities favors the explicitly evaluation of current metacommuity theories. On one side, a classical interpretation of the relationships between environmental conditions and the composition of fish communities is closely linked to Niche theory, predicting that the resources and conditions of a given environment dictate the distribution of species over space and time [1]. Studies of relationships between species composition, environmental conditions (e.g. water temperature, dissolved oxygen content, substrate type) and resources (e.g. places of refuge, food) have had an important impact on our understanding of how those systems work [1]. Considering the four general models currently used to explain metacommunity patterns [2], Species-Sorting assumes that the environment gradient is the only factor structuring the communities [3–7]. On the other hand, the explicitly spatial strucuture of riverine systems [8] and the existence of some complex models of linkage among aquatic habitats within basin (e.g. river pulse and marginal lakes), favors the view that dispersal within the metacommunity may account for a signification portion of the explanation of species composition patterns [7]. This later explanation is usually refered as a neutral framework [2]. Even though both factors may interact, we are still looking for a set of more general predictions from which it is expected that each of the models can be better applied.

The effect of environmental conditions such as water velocity, dissolved oxygen and water temperature successifully explain common patterns observed on fish communities of tropical streams [9–14]. Nevertheless, the choice of environmental descriptors and the quality of its statistical description is subject to doubts. Often these conditions are measured as a single value [12,15]. In contrast, the use of repeated measurements [e.g.: 11,16–18] may favor a better description of the natural stochasticity of environmental conditions in streams. Better designs for environmental conditions measurement may account for environmental heterogeneity of the stream system, which is recognized a long time [19–21], but frequently neglected. Environmental heterogeneity can have important effects on biological diversity [22,23], suggesting that variation of in-stream conditions is a more important predictor of fish species diversity [e.g.: 19–21], and others aquatics communities than average values [e.g.: 24,25]. For example, the relationship between aquatic macroinvertebrate diversity and the heterogeneity of substrates in streams [26–28].

Dispersal-related process may also exert some challenges related to data acquisition and interpretation The Neutral Theory [29] proposes that all individuals of the same trophic guild, have the same ability to compete (regardless of resources and present conditions at the place), thus having constant growth in any environment. In such scenario the author proposes that dispersal is the main source of increase local community diversity at an ecological time framework. Thus, it is expected that the spatial component (distance between sampling sites) or geographical barriers (physical obstacles to migration between locations) are the only mechanisms responsible for structuring the community. In the studies of stream ecology, we observe the frequent use of linear distance as a measurement of connectivity. However this distance may be not informative, since strict aquatic organisms (such as fish) can only move through the hydrographic network. Connectivity measures based on the stream network is beginning to be used in the interpretation of patterns on estuarine fish communities [e.g.: 30] and other aquatic organisms [e.g.: 28,31,32]. The main problem of the use of linear distances is the comparison of streams present in different hydrographic units. Usually, streams in the same basin (or sub-basins) are more similar than streams in different basins even if streams in different basins are geographically closer (linear distance) than streams within the same basin (hydrographic distance) [28].

There is no simple dichotomy between communities structured by Niche processes and neutral processes in nature [3], as many natural communities are structured both by local conditions and resources (Niche-related processes), and species’ dispersion abilities [3,33]. The combined effects of different processes on communities are united under the theory of Mass-effects [2], which describes meta-community processes as the product of both environmental gradients and the dispersal ability of the species. The quantification of the relative importance of niche and neutral processes can be evaluated using variance partitioning techniques [3,20,34]. These techniques allow to quantifying the percentage explanation related only to environmental conditions and resources, only to space, and to the interaction between the two sets [35]. Obviouly, the choice of descriptive variables for both sets – environment and space – may have a determinant effect on the results and on the interpretation of how both explanations interact.

The evaluation of how space and environmental variables are related to community composition allow to a more explicity analysis of the now classic mechanisms for metacommunity patters developed by Leibold et al [2]. Communities that relate solely to the environmental gradient would be structured by the Species sorting mechanism, whereas communities that relate only to space are structured by Neutral or Patch Dynamics, and the interaction between space and the environmental gradient is taken as communities where Mass-Effect processes are predominant [2,3]. In the present study, our goals were: (i) identify the best representation of local conditions (average values or variance of conditions), and (ii) evaluate the importance of linear and hydrographic distance in Neutral processes; (iii) evaluate the relative explanatory power of environmental and space variables in order to determine wich of the four models proposed by Leibold account for observed patterns in Brazilian Cerrado fish communities.

## Materials and Methods

To test the hypotheses we used a database composed of 76 streams of three distinct hydrographic regions of Brazil: i) Araguaia-Tocantins; Ii) Paraná; and iii) São Francisco, with all streams are located in the Cerrado biome (Figure 1). All sampled streams are first-to third-order, georeferenced and the ichthyofauna was sampled by trawl or electric fishing at least 50 m from the channel. The local conditions measured were turbidity, conductivity, pH, dissolved oxygen, velocity, width and depth in at least three points of each stream.

**Figure 01.**
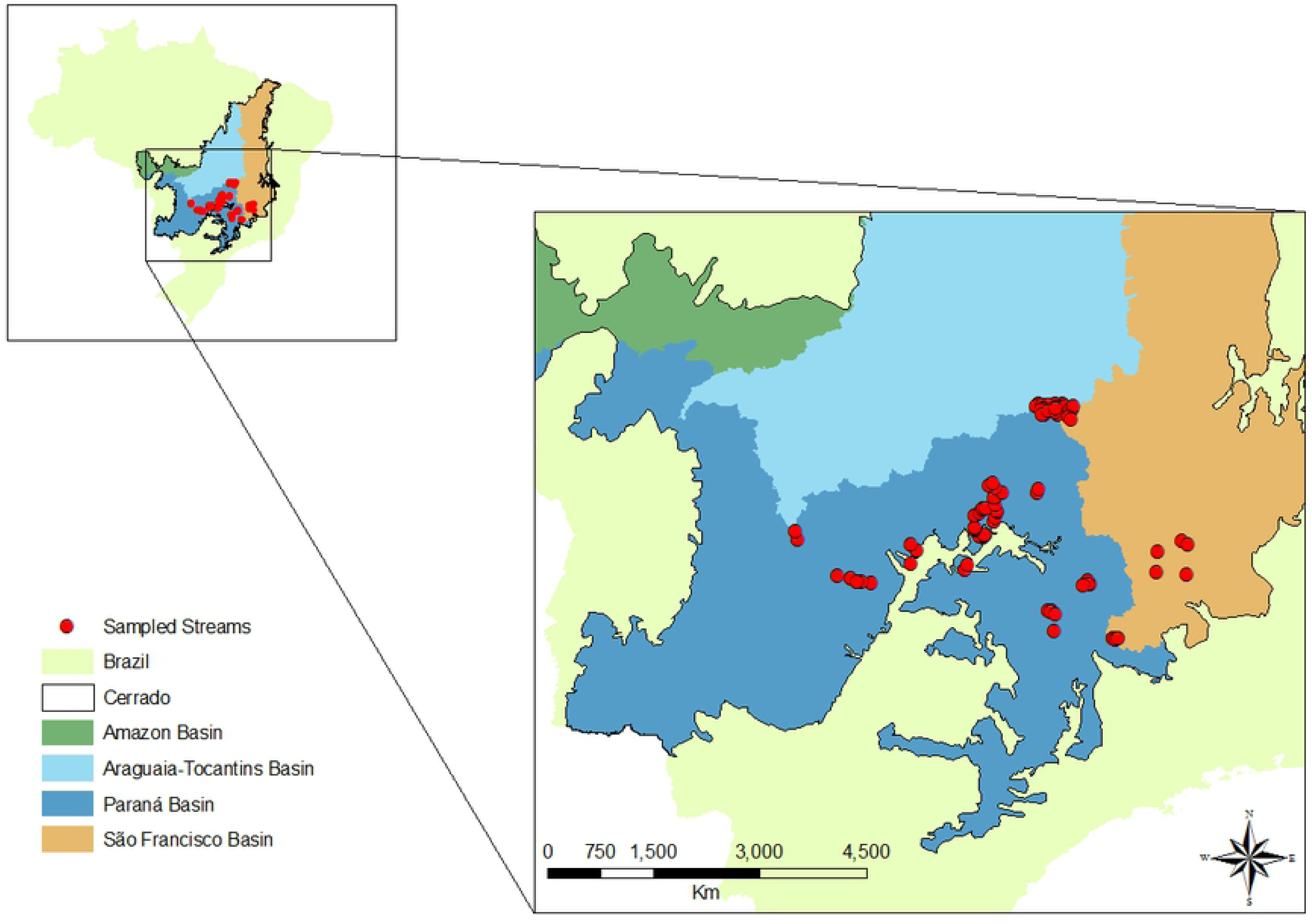
Location of the 76 streams (points) used in the work in three hydrographic regions in the Brazilian Cerrado.

As the points sampled are geographically aggregated, the autocorrelation of the data was tested through the Moran’s Index. Spatial maps of suitable vectors are considered the best way to control autocorrelation and are good representatives of the spatial structure present in the data [36]. The eigenvectors of the first eigenvalues are related to the local structuring of communities, species with small dispersion skills. In order to perform spatial eigenvectors maps is necessary to know the relationship between all pairs of points present in the analysis, known as the Weight (W) Matrix [37]. In this way we define four W: i) Global W (Appendix 1 - S1 - GW), defined by the linear distance between all streams (Appendix 2); ii) Local W (Appendix 1 - S1 - LW), defined by the linear distance between streams present in the same hydrographic unit. Streams in different units had no interaction and connectivity values equal to zero (Appendix 2); iii) Water W (Appendix 1 - S1 - WW), defined by the hydrographic distance. The same way as in Local W, streams in different units had no interaction and connectivity values equal to zero (Appendix 2); and iv) W Hydalt (Appendix 1 - S1 - HW), defined by the hydrographic distance between the points weighted by the slope (Appendix 2).

We measured ecological diversity using two distinct metrics: (i) species richness and (ii) beta diversity. Species richness was defined as the number of species present at the site of interest. The beta diversity was calculated according to the procedure described by Baselga [38], which defines beta diversity as the Sorensen dissimilarity index. Diversity indices were calculated for both the total fish community and for different feeding guilds (i.e. detritivores, insectivores and omnivores).We used generalized linear models (GLM) to identify the best way to represent environmental conditions for streams fish communities. We compared models of community diversity indices and the average of the environmental conditions with a second model including the average together with the standard deviations of environmental conditions as predictor variables. This procedure was adopted to identify the best way to describe environment conditions, the average or variance. The GLM models was performed using all W matrices (Table 1) to identify the best means to represent neutral processes (linear or hydrographic distances). To avoid multicollinearity, a Principal Component Analysis (PCA) was performed with the average values, and a second PCA with the average values together with the standard deviation values of the descriptors, the first two axes of explanation of the PCA being used as predictor variables on the models. After identifying the best way to represent environmental conditions (mean or mean with standard deviation), GLMs were performed for each trophic guild and all W matrices. This procedure was used to identify if results found for the whole community were equivalent to those found for individual feeding guilds.

**Table 1.** Models used to determine the best set of descriptors of environmental conditions and spatial structure, considering the richness and beta diversity of ichthyofauna.

| Diversity Metric | W Matrix | Environmental Condition |
| --- | --- | --- |
| Richness | Global W | Average<br>Average plus Standard Deviation |
|  | Local W | Average<br>Average plus Standard Deviation |
|  | Water W | Average<br>Average plus Standard Deviation |
|  | Hydalt W | Average<br>Average plus Standard Deviation |
| Beta Diversity | Global W | Average<br>Average plus Standard Deviation |
|  | Local W | Average<br>Average plus Standard Deviation |
|  | Water W | Average<br>Average plus Standard Deviation |
|  | Hydalt W | Average<br>Average plus Standard Deviation |

All analyses were performed in the Spatial Analysis for Macroecology - SAM [39]. For all models, we compute the Moran’s I and the Akaike Information Criterion (AIC) and the Variation of the AIC (ΔAIC). To test relative effects of different metacommunity processes, variance was partitioned into: i) environmental conditions; ii) spatial processes; iii) shared variance between the environmental and space; and iv) not explained.

## Results

The richness and beta diversity of ichthyofauna had a significant spatial structure with values of Moran’s I higher than 0.1 (Table 2). Richness of omnivores showed the greatest degree of spatial structure (Moran’s I = 0.316, p <0.001), and the richness of detritivores the lowest (Moran’s I = 0.120, P = 0.010). The beta diversity of the detritivores, insectivores and omnivores presented non-significant effects of spatial autocorrelation.

**Table 2.** Autocorrelation index of Moran (Moran’s I) performed for richness and beta diversity for all community and by guilds. Moran’s I - Autocorrelation index of Moran; p - Type one error probability; I (max) - The highest value of Moran’s I to the variable; I/I(max) - The Moran’s I corrected by maximum value.

| Diversity Metric | Ichthyofauna | Moran's I | $p$ | I (max) | I/I(max) |
| --- | --- | --- | --- | --- | --- |
| Richness | All | 0.234 | <0.001 | 0.737 | 0.318 |
|  | Detritivorous | 0.120 | 0.010 | 0.610 | 0.198 |
|  | Insectivorous | 0.180 | <0.001 | 0.628 | 0.287 |
|  | Omnivorous | 0.316 | <0.001 | 0.807 | 0.392 |
| Beta Diversity | All | 0.208 | <0.001 | 0.758 | 0.274 |
|  | Detritivorous | 0.052 | 0.211 | 0.680 | 0.076 |
|  | Insectivorous | 0.020 | 0.517 | 0.551 | 0.037 |
|  | Omnivorous | -0.039 | 0.618 | 0.748 | -0.052 |

The PCA performed with average values of the environmental conditions explained 33.26% of variation in fish diversity in the first axis, and 18.79% in the second axis, and a total of 52.05% across both axes (Appendix 3). The pattern found by the ranking was: (i) Conductivity, Ph, Dissolved Oxygen and Channel Width positively related to the first axis; (ii) Turbidity and Current Speed positively to the second axis; and (iii), Channel Depth negatively to the second axis (Appendix 3). The PCA performed with both average values and standard deviation of environmental conditions explained 21.02% of variance in the first axis and 16.66% in the second axis (Appendix 3). The pattern found by the ranking was: (i) Standard deviation of the channel width, Standard deviation and Average Turbidity and Current Speed positively related to the first axis; (ii) Standard Deviation and Average Depth of the channel and the Standard Deviation of Ph, Conductivity and Oxygen dissolved negatively to the first axis; and Average Channel Width, Ph, Dissolved Oxygen and Conductivity and the Standard Deviation of Water Temperature positively to the second axis (Appendix 3).

Best models of richness and beta diversity of fish species from included average and standard deviation of the local conditions and the spatial eigenvalues maps performed from Local W (Appendix 3). The richness model had a high r-squared value (r^2^ = 0.623), with 56.30% of variance explained by environmental conditions, 6% by spatial maps and 0.3% by interactions between niche and neutral effects (Appendix 3). Beta diversity had an even higher r-square value (r^2^ = 0.758), with 64.40% of variance explained by environmental conditions, 6.70% by spatial maps, and 4.80% by interactions between environmental and spatial processes (Appendix 3).

When the community was partitioned into trophic guilds, richness and beta diversity of insectivorous and omnivorous species presented the spatial eigenvalues maps, performed from the W Local matrix, as the best representation of spatial processes (Appendix 3). However, for detritivores, the W Hydalt model was the best representation of spatial processes (Appendix 3).

## Discussion

In the analyses of richness and beta diversity, we found environmental conditions captured most of the variation in fish species diversity, suggesting that environmental conditions were satisfactorily represented using average values, and the spatial component contributed little to structuring communities. Interpretation of spatial processes and association with dispersion, made using spatial eigenvalue maps, has to be done carefully. Spatial eigenvalue maps bring not only information on spatial structuring but also environmental conditions [40,41].

As for beta diversity in different guilds, especially insectivores and the omnivores, showed little relation with environment conditions and a greater correlation with spatial processes. This result may be explain by the spatial component representing collinear unmeasured environmental conditions rather than effects of space itself, or that insectivorous and omnivorous fish have greater dispersion abilities. A higher dispersal capacity can hide effects of environmental gradients, as species can rapidly colonize sites that have unfavorable conditions, and suppress effects of local extinction [42]. If this was the case for insectivorous and omnivorous species, Mass Effects may be the predominant mechanism structuring their communities. In contrast, detritivores and the total community were structured by Species Sorting effects, or an interaction between Mass Effects and Species Sorting [3], as these communities showed a strong relationship with environmental conditions and little correlation with space.

Although the Local W model was the best representation of space for the entire ichthyofauna community, we observed a weak relationship between fish communities (richness and total beta diversity) and spatial components. This weak association may have occurred due two factors: (i) The ability of fish to actively select habitats would be more important than dispersion, indicating the Species-Sorting mechanism of metacommunity structure [2]. Species Sorting is considered the main mechanism responsible for structuring natural meta communities [3,4,6,7]. On the other hand, (ii) the “path” along which fish species have dispersed is environmentally unsuitable, restricting their dispersion [42]. Although the characteristics of streams were controlled in order to be representative of natural conditions, the connection between these points was not controlled. Thus, it may be that sites, although preserved, are connected by non-suitable drainage that act as a barrier and limit the dispersion of ichthyofauna [42].

Linear distances without considering hydrographic units fail to represent spatial process in stream fishes communities. The use of the linear distance is a simplistic and insufficient means measuring spatial processes in aquatic systems [7]. When including the geographic barriers into the linear distance procedure (called Local W here), the linear distance was able to provide a good representation of the geographic patterns of this fish community. The performance of Local W is better than the dendritic distance (W Water), shown previously as the best means to represent spatial processes in aquatic systems. Using Local W, we were able to identify a simple and robust way of representing spatial processes considering the physical barriers separating communities. This result was consistent across all components of the community (*i.e.* overall community or discrete feeding guilds) and ecological descriptors (*i.e.* species richness or beta diversity.

A second feature of our results is related to the dispersion of detritivorous fish, which cannot be represented using simple euclidian distances between points, considering or not considering geographic barriers, or hydrological distances. The best way to represent space process in this guild was with the application of Hydalt W. These models consider the dendritic distance between points (and barriers), and the direction of the flow. The application of connectivity models associated with flow direction is advocated by some authors as being the most suitable for aquatic organisms [43,44]. However, with the analysis of our data, we observed that this approach, is more complex than the euclidian distance (considering barriers - Local W), and was only necessary for detritivorous fish.

The low dispersion present in the ichthyofauna of streams is evidenced in the negative relation between communities and spatial filters. Considering that small eigenvalues are related to local structuring and higher eigenvalues to larger scale structuring [37,45], we interpret that fishes from stream are dispersing across small scales, as richness and beta diversity were negatively related to the filters. This result supports the idea of preserved spots connected by altered “paths”. Thus, species with low dispersal ability tend to express spatial structure more clearly than species with higher dispersal ability [33,46].

Species-Sorting predicts a correlation of metacommunity structure only with environmental gradients [2]. The interaction of space (dispersion) and environment in the structuring of metacommunities is attributed solely to the Mass Effects model [2]. However is it not possible to separate these two mechanisms, and metacommunities that are related to space and environment are understood as interaction between Species-Sorting and Mass Effects [3,47–49]. This interaction was found in 29% (46) of the communities analyzed by Cottenie [3] and also found for Amazonian fish [7,47–49]. However the ichthyofauna show a greater correlation with environmental conditions than spatial processes, suggesting that Species Sorting is the key mechanism in structuring fish communities (37% of the metacommunity analyzed by Cottenie [3] show this result). This result does not rule out the occurrence of dispersion [3,7,47–49], but reinforces the idea that dispersion occurs locally and weakly (negative relation with the spatial eigenvectors maps).

Species Sorting is related to many freshwater organisms, such as macroinvertebrates [42,50], snails [51] and bacteria [4]. The power of the Species Sorting mechanism in communities at large scales was found showing that there was no variation in the metacommunities between three basins studied, but a continuous variation according to an environmental gradient [52]. Dispersion has little influence on metacommunity patterns at large scales [42,50]. In fact, dispersion decrease at large scales as the same way that environmental gradient increase the correlation with metacommunity structure [42,46]. In addition, fish can only migrate using drainage and some species tend not to migrate or are prevented from migrating due to physical barriers such waterfalls or dams. Furthermore, species that actively disperse tend to select the environment in which they will settle, further reducing effects of spatial structure [42].

The variation partitioning of fish communities between environmental conditions and geographic space tend to associate these sets of variables with niche (i.e. environmental conditions) or neutral (i.e. geographic distances) theory [53]. When communities are more correlated with environmental conditions than geographic distance, the communities are structured by niche theory. On the other hand, when the geographic distance are more correlated than environment, the discussion is focused on neutral theory. This dichotomous view is overly simplistic and does not reflect complexities of the multiple mechanisms that concurrently structure natural communities. Communities are not the effect of only one of these two theories, but the interaction between them [54]. More recent analyses [3,4,42,46–49,55] demonstrate the association of the community with the mechanisms proposed by Leiboldet et al [2]. In this case, the relationship between communities and environmental conditions are related to the Species Sorting mechanism, the relationship between community and space to the neutral or Patch Dynamic mechanisms, and the interaction between space and conditions related to the Mass Effect mechanism [3]. However, the dissociation of Mass Effects and Species Sorting is not trivial, since the dispersion limitation in communities within Species-Sorting can produce a pattern with relationship between space and environmental conditions, a Mass Effect dynamic [3].

So we conclude that the ichthyofauna of Cerrado streams are structured by an interaction between Mass Effects and Species Sorting mechanisms. Only the beta diversity insectivores and omnivores differed, with stronger effects of space, suggesting an influence of neutral or Pach Dynamic models. Finally, we found that the linear distance measures that take into account the physical barriers (i.e. Local W) are the best representations of spatial patterns in fish communities. Although, analyses of detritivores required the addition of flow direction in connectivity models (Hydalt W). Regarding the environmental conditions, it is necessary that these are represented by average and some metric that measures their variation to best evaluate the degree of environmental heterogeneity.

## Acknowledgements

We would like to thank the lab staff at the Aquatic Ecology Laboratory of the UFC. TBV would like to thank Coordenação de Aperfeiçoamento de Pessoal de Nível Superior (CAPES) for the grant provided during his doctoral degree. Part of these study was funded by CNPq/PPBIO (agreement # 457463/2012-0) and CNPq/ICMBio (agreement # 552009/2011-3), CNPq grant (Process 471283/2006-1), The International Fund for Ecological Research, grant no. 00543 and Aperfeiçoamento de Pessoal de Nível Superior - Brasil (CAPES) Finance Code 001

## Supporting information captions

Appendix 01 - Weight (W) Matrix used in the analysis. i) Global W (S1 – GW); ii) Local W (S1 - LW); iii) Water W (S1 - WW); iv) W Hydalt (S1 - HW).

Appendix 02 – Procedures performed to calculate all W matrices

Appendix 03 – Results from PCA analyses and GLM procedures performed with communities, environmental conditions and space processes.

